# The Lysosome Surface is an Unappreciated Hub for Vasopressin V2 Receptor Signaling

**DOI:** 10.1101/2025.10.03.680346

**Authors:** Emmanuel Flores-Espinoza, Hyunggu Hahn, Alex R.B. Thomsen

**Affiliations:** Department of Molecular Pathobiology, New York University College of Dentistry, New York, NY 10010, USA; NYU Pain Research Center, New York University College of Dentistry, New York, NY 10010, USA

## Abstract

G protein-coupled receptors (GPCRs) have traditionally been understood to signal through heterotrimeric G proteins exclusively from the cell surface followed by β-arrestin (βarr)-mediated desensitization and receptor internalization into endosomes. However, this view has evolved significantly with growing evidence showing that some GPCRs continue to signal from endosomes after their internalization as well as from other intracellular organelles. The vasopressin V2 receptor (V2R) exemplifies this paradigm shift as it promotes robust endosomal G protein before being sorted to lysosomes for degradation. Intriguingly, recent observations suggest that the lysosomal surface itself holds a substantial pool of heterotrimeric G proteins, raising the possibility that GPCRs such as the V2R may stimulate signaling from this subcellular region. To investigate this, we here employed a NanoBiT bystander approach to track intracellular V2R trafficking and transducer activation in real-time. Our results show that activated V2R is trafficked relatively fast to lysosomes where it retains the ability to couple to both G proteins and βarrs. Applying nanobody/intrabody biosensors, we further demonstrated that the V2R activates endogenous G proteins and βarrs at the lysosomal surface and that inhibition of V2R translocation to endolysosomal compartments blunts its ability to stimulate G protein signaling. Together, these findings suggest that the lysosomal surface serves as an unappreciated hub for signaling by some GPCRs before they eventually are engulfed into the lysosomal lumen for degradation.

**One-sentence summary:** Upon activation, the vasopressin V2 receptor is internalized and sorted to the lysosomal membrane, where it activates G proteins and β-arrestins.

## Introduction

The vasopressin V2 receptor (V2R) belongs to the superfamily of G protein-coupled receptor (GPCRs), cell surface proteins that regulate virtually all physiological processes and are important drug targets. The V2R is mainly expressed at the basolateral membrane of the kidney collecting duct principal cells and is activated by arginine vasopressin (AVP), a nine-amino acid hormone that is secreted by the posterior pituitary during hypovolemic conditions (*1, 2*) Once activated, V2R stimulates heterotrimeric G_s_, which triggers cAMP production and activates protein kinase A (PKA) (*3*). Activated PKA then phosphorylates aquaporin-2 (AQP2) located at intracellular vesicles, which leads to insertion of these vesicles into the apical membrane (*4*). In addition, activity of the sodium channel ENaC on the apical membrane is enhanced leading to sodium uptake by collecting duct cells. The overall effect of these actions enhances renal water reabsorption and the concentration of urine, both of which help maintain water homeostasis (*1, 5*).

The V2R contains clusters of serine/threonine phosphorylation sites in its carboxy (C)-terminal tail and forms highly stable interactions with β-arrestin1/2 (βarr1/2) upon activation and phosphorylation (*6–8*). These stable V2R–βarr complexes promote profound receptor internalization into early endosomes (*9*), from where the receptor eventually is sorted to lysosomes for degradation (*10*). This stable interaction with βarrs and trafficking pattern are characteristic of a subset of GPCRs, referred to as class B, that harbor phosphorylation site clusters in their C-terminal tail and/or third intracellular loop (*9*). In contrast, class A receptors, such as the β2-adrenergic receptor (β2AR), only contain few individual serine/threonine phosphorylation sites and therefore associate more transiently with βarrs, resulting in rapid recycling of internalized receptors back to the cell surface (*7*). Interestingly, we and others have demonstrated that class B GPCRs such as the V2R can form stable complexes with βarrs exclusively via the phosphorylated C-terminal tail (*11–13*). As βarrs in this ‘tail’ conformation does not occupy the G protein binding site within the receptor core, these GPCRs continue to activate G protein while being internalized into endosomes in complex with βarrs (*11, 14*). Interestingly, sustained G_s_ protein signaling by internalized V2R has been linked to apical membrane transport of ENaC and enhanced phosphorylation of AQP2 in kidney collecting duct cells (*15*).

Although endosomal G protein signaling appears to be most persistent for class B GPCRs, increasing amount of evidence suggests that other receptors remain signaling competent after their internalization (*16–21*). In fact, intracellular GPCR activity is not just be confined to endosomes but has also been observed at other organelles including membranes of the endoplasmic reticulum (*22*), trans-Golgi network (*23, 24*), mitochondria (*25, 26*), and nuclear envelope (*27, 28*). Moreover, GPCR activity at specific organelles has been linked to distinct cellular functions compared to signaling at the plasma membrane, highlighting the importance of understanding the spatiotemporal regulation of GPCR signaling (*29–32*). An organelle that remains relatively unexplored in the context of compartmentalized GPCR signaling is the lysosome. Lysosomes are acidic organelles (pH ∼4.5) that contain hydrolases and serve as the final destination for GPCRs and other membrane proteins where they ultimately are degraded (*33, 34*). Based on this function, it would seem unlikely that lysosome could act as a signaling hub for internalized GPCRs. However, a recent study demonstrated that a significant pool of endogenous heterotrimeric G protein exists at the lysosomal surface raising the possibility that G protein signaling may occur from this subcellular location (*35*). In the case of V2R, its trafficking to lysosomes occurs after prolonged stimulation with AVP (*36*), and although the receptor is degraded here, this process is slow. Therefore, we here used a NanoBiT bystander approach to investigate if the V2R retains its ability to activate G proteins and βarrs at the lysosomal surface before eventually being engulfed and degraded.

## Results

### V2R trafficking to lysosomes depends on βarr1/2

The trafficking of V2R from the cell surface to early endosomes and lysosomes was followed in real-time using a NanoBiT molecular bystander approach. For this purpose, the small BiT component of the NanoBiT system was fused to the C-terminal tail of the V2R (V2R-SmBiT) while the large BiT (LgBiT) part was linked to subcellular markers of the plasma membrane (CAAX-LgBiT), early endosome (FYVE-LgBiT), or lysosomes (LAMP1-LgBiT) (Fig. 1A). When co-expressed in HEK293 cells, close proximity of V2R-SmBiT and subcellular marker-LgBiT results in the formation of a functional nanoluciferase enzyme that can be used to report on the presence of the receptor at the different subcellular compartments. As the LAMP1-LgBiT construct has not been used previously, we expressed it together with the commercially available CellLight™ Lysosomes-RFP marker in HEK293 cells. In addition to the LgBiT part, the LAMP1-LgBiT construct also contains a HA-tag, which allowed us to localize it in cells by immunostaining using an anti-HA primary antibody. Imaging cells expressing these two constructs by confocal microscopy revealed that they almost perfectly colocalize (Supplementary Fig. S1A-B). This observation validates that the LAMP1-LgBiT can be used as a lysosomal marker for NanoBiT experiments.

**Figure 1.**
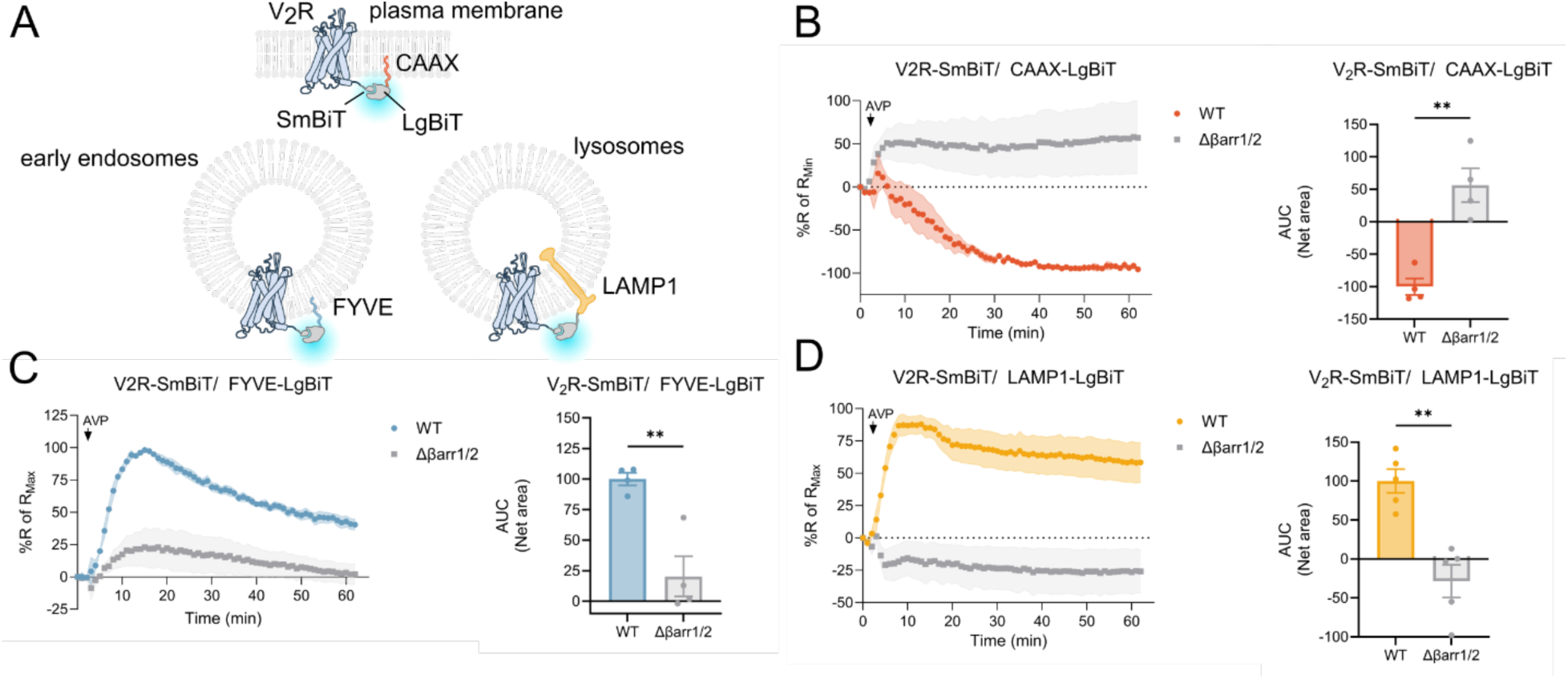
Intracellular V2R trafficking and its dependency on βarr1 and receptor C-tail. (**A**) Schematic illustration showing how SmBiT and LgBiT-conjugated NanoBiT constructs were used to follow the trafficking of V2R-SmBiT from the plasma membrane (CAAX-LgBiT used as marker) to early endosomes (FYVE-LgBiT used as marker) and lysosomes (LAMP1-LgBiT used as marker). (**B-D**) Real-time changes in luminescence signal generated between V2R-SmBiT and CAAX-LgBiT (**B**), FYVE-LgBiT (**C**), or LAMP1-LgBiT (**D**) upon AVP-stimulation (100 nM). Responses were normalized to the maximal change in luminescence. Experiments conducted in both HEK293 parental cells (WT; colored) and HEK293 Δβarr1/2 cells (gray) are displayed. Area under the curve (AUC) values were used to calculate the total receptor trafficking response (bar graphs). Data represent the mean ± SEM from N=4 independent experiments. Unpaired t-test with two-tail p-value tests were applied to determine the statistical differences between the distinct treatments (**p < 0.01).

Upon AVP-stimulation of V2R-SmBiT/CAAX-LgBiT expressing cells, we observed a fast spike in signal within the first 2 minutes followed by a robust reduction in the luminescence signal. This indicates that V2R-activation initially leads to a very fast and surprising increase in receptors at the cell surface followed by robust and a long-lasting internalization phase (Fig. 1B). Simultaneously, the signal reporting on V2R presence in early endosomes increased steadily with a peak around ∼12 minutes after AVP challenge in V2R-SmBiT/FYVE-LgBiT expressing cells (Fig. 1C). This initial phase of V2R-enrichment at early endosomes was followed by a slow decrease throughout the experiment. Interestingly, when V2R-SmBiT/LAMP1-LgBiT expressing cells were treated with AVP, receptor trafficking to lysosomes was confirmed but with very similar initial kinetics as V2R trafficking to early endosomes (Fig. 1D). In contrast to V2R trafficking to early endosomes, the V2R at lysosomes remained elevated throughout the experiment (Fig. 1D). The presence of internalized V2R in early endosomes and lysosomes after 30 minutes of AVP challenge was further confirmed by confocal microscopy in HEK293 cells expressing a SNAP-tagged V2R (Supplementary Fig. S1C-D).

Under all three NanoBiT expression conditions, the V2R internalization and translocation to early endosomes/lysosomes signals were blunted when the experiments were performed in βarr1/2 double knockout (DKO) HEK293 cells (Δβarr1/2) (Fig. 1B-D). These results suggest that V2R internalization and intracellular trafficking is mainly driven by complex formation with βarrs as expected. A minor, yet significant, increase in translocation of V2R-SmBiT to FYVE-LgBiT, but not to LAMP1-LgBiT, was observed in response to AVP stimulation in HEK293 Δβarr1/2 cells (Fig. 1C-D). However, similar βarr-independent V2R translocation to early endosomes has previously been reported (*11*).

### βarr1 translocates to the lysosomal surface in response to prolonged V2R activation

βarrs are known to play a major role in facilitating GPCR internalization and intracellular trafficking (*37*). In fact, they associate particularly well with active V2R via phosphorylated cluster sites within the receptor C-terminal tail, which leads to robust receptor internalization into endosomes followed by sorting to the lysosome similar to other Class B GPCRs (*9, 36*). Whether βarrs stays associated with the V2R during lysosomal sorting process is poorly understood. Therefore, we decided to investigate how βarr1 translocate to different subcellular compartments including the plasma membrane, early endosomes and lysosomes upon V2R activation (Fig. 2A). For this purpose, HEK293 cells were co-transfected with V2R, βarr1-SmBiT and one of the three compartment markers: CAAX-LgBiT (Fig. 2B), FYVE-LgBiT (Fig. 2C), or LAMP1-LgBiT (Fig. 2D). Upon AVP-stimulation, βarr1-SmBiT was recruited robustly and fast to the plasma membrane CAAX-LgBiT marker (1-2 min) followed by gradual removal from this subcellular location (Fig. 2B). AVP-provoked βarr1-SmBiT translocation to the early endosomal marker, FYVE-LgBiT, was also robust but slower than its recruitment to the cell surface (∼12 min, Fig. 2C). Interestingly, translocation of βarr1-SmBiT to the lysosomal marker LAMP1-LgBiT in response to AVP challenge displayed similar kinetics as the recruitment of βarr1 to early endosomes (Fig. 2D). However, V2R-βarr1 association at lysosomes was more persistent than at early endosomes (Fig. 2D). Under all three NanoBiT expression conditions, no AVP-stimulated translocation of βarr1 was observed in HEK293 cells that do not express the V2R (Fig. 2B-D). Together, these results suggest that after V2R-βarr1 complexes have been internalized into early endosomes, they are quickly sorted to the surface of lysosomes where they stay for some time before being engulfed into the lumen and degraded.

**Figure 2.**
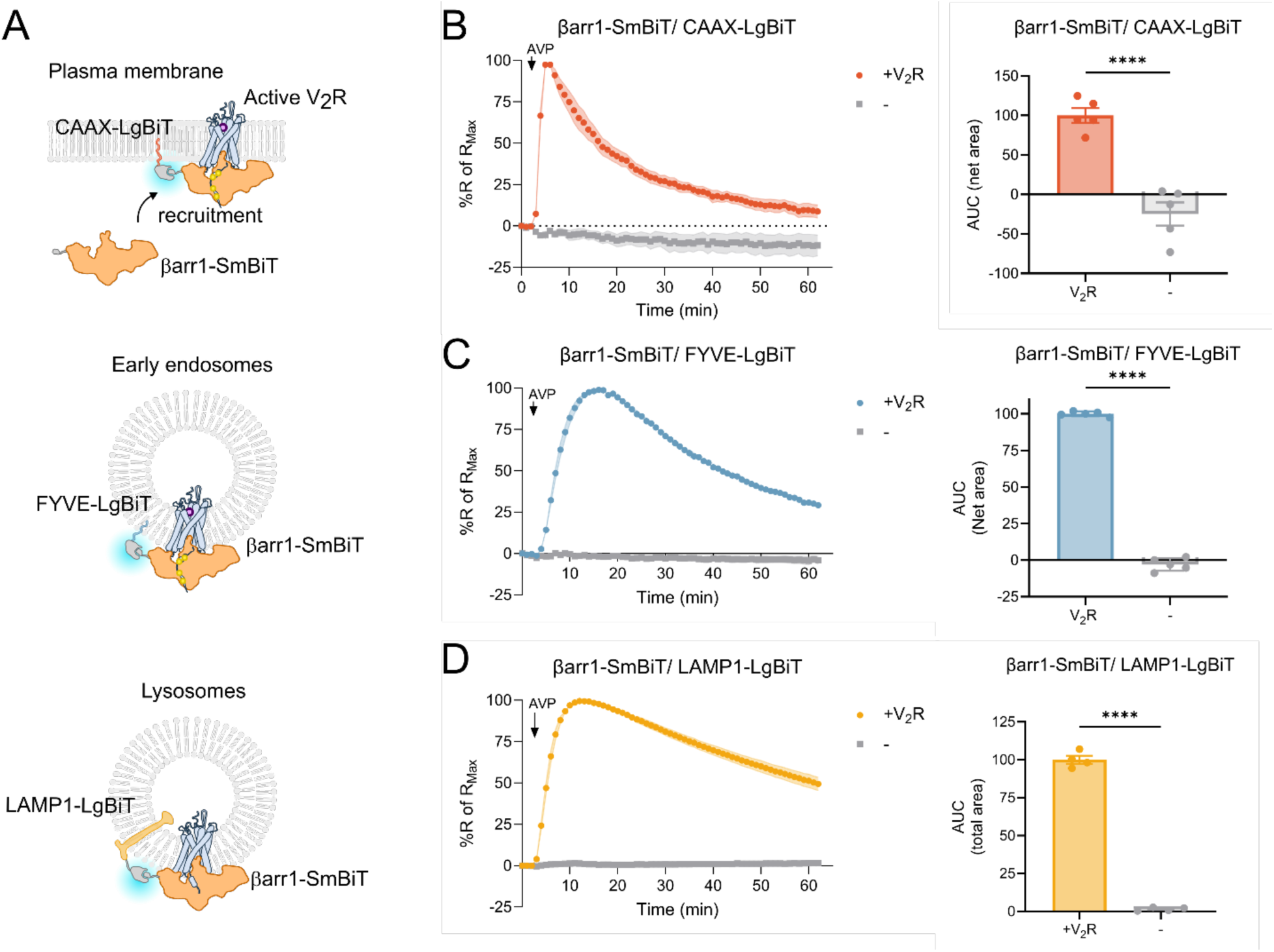
V2R-stimulated βarr1 translocation to the plasma membrane, early endosome, and lysosome. **(A**) Experimental design to measure βarr1-SmBiT recruitment to different cell compartments using the NanoBiT assay. (**B-D**) Real-time changes in luminescence signal generated between βarr1-SmBiT with CAAX-LgBiT (**B**), FYVE-LgBiT (**C**), and LAMP1-LgBiT (**D**) in response to AVP (100 nM) stimulation. Responses were normalized to the maximal change in luminescence. The experiments were conducted in V2R-expresing HEK293 cells (colored curves) and non-transfected HEK293 cells (gray curves). AUC values were used to calculate the total βarr1 translocation response (bar graphs). Data represent the mean ± SEM from N=4-5 independent experiments. Unpaired t-test with two-tail p-value tests were applied to determine the statistical differences between the distinct treatments (****p < 0.0001).

### AVP-stimulated and internalized V2R maintains its ability to couple to G proteins at lysosomes

When activated, V2R couples principally to G_s_, and often to G_q/11_, even while being bound to and internalized by βarrs in the tail conformation (*11, 12, 14*). As the V2R–βarr1 complex is trafficked to first early endosomes and then sorted to lysosomes, we asked whether the V2R maintain its ability to couple to G proteins at these subcellular compartments. Recently, it was demonstrated that a large pool of endogenous heterotrimeric G proteins is found at the lysosomal surface (*35*), and thus, we were particularly interested in whether the V2R can activate this pool of G proteins.

To determine if the V2R maintain its ability to couple to G protein at the membrane of different subcellular compartments, we used a strategy based on mini-G_s_ and mini-G_s/q_ proteins (denoted as mGs and mGq here), which act as engineered surrogates of the G_s_ and G_q/11_ coupling to GPCRs, respectively (*38, 39*). One of the key differences in mG proteins from corresponding wild-type Gα subunits is a truncation of the amino (N)-terminal that prevents membrane anchoring, resulting in their cytosolic localization under resting conditions (*40*). Upon agonist stimulation, mGs- or mGq-SmBiT translocate from the cytosol to specific subcellular membranes where the activated receptor reside, which can be quantified effectively by the NanoBiT system using the LgBiT-attached compartmental markers (Fig. 3A). When either mGs-SmBiT or mGq-SmBiT were co-expressed with plasma membrane marker CAAX-LgBiT in V2R-expressing HEK293 cells, AVP-stimulation led to fast translocation of both mG proteins to the plasma membrane, followed by gradual dissociation (Fig. 3B and 3E). No recruitment of either mGs-SmBiT or mGq-SmBiT was detected upon AVP challenge in HEK293 cells that do not express the V2R (Fig. 3B and 3E). Curiously, AVP-stimulation of HEK293-V2R cells, but not non-V2R cells, expressing the early endosomal marker, FYVE-LgBiT, caused an initial fast drop in the luminescence signal with both mGs-SmBiT and mGq-SmBiT. This initial drop in luminescence was followed by a robust increase that remained elevated throughout the rest of the experiment (Fig. 3C and 3F). Similar translocation of mGs/mGq to the lysosomal marker LAMP1-LgBiT was observed upon V2R stimulation, but without the initial drop in the luminescence signal (Fig. 3D and 3G). As with βarr1 recruitment, the translocation of mGs/mGq to early endosomes or lysosomes appeared to have somewhat similar kinetics but remained more sustained elevated in the lysosome. For unclear reasons, mG_q_ translocation to both early endosomes and lysosomes was slower than with mGs. Furthermore, AVP-induced recruitment of mGs-SmBiT to LAMP1-LgBiT was completely blunted in HEK293 Δβarr1/2 cells (Supplementary Fig. S2A-B), indicating that the translocation of mGs to lysosomes depends on βarrs (presumably as βarrs traffic the V2R to this subcellular compartment). Finally, no response to AVP challenge was observed in HEK293 cells that do not express V2R (Fig. 3D and 3G). Together, these results suggest that the V2R is fully capable of interacting with G proteins during its trafficking to early endosomes and lysosomes.

**Figure 3.**
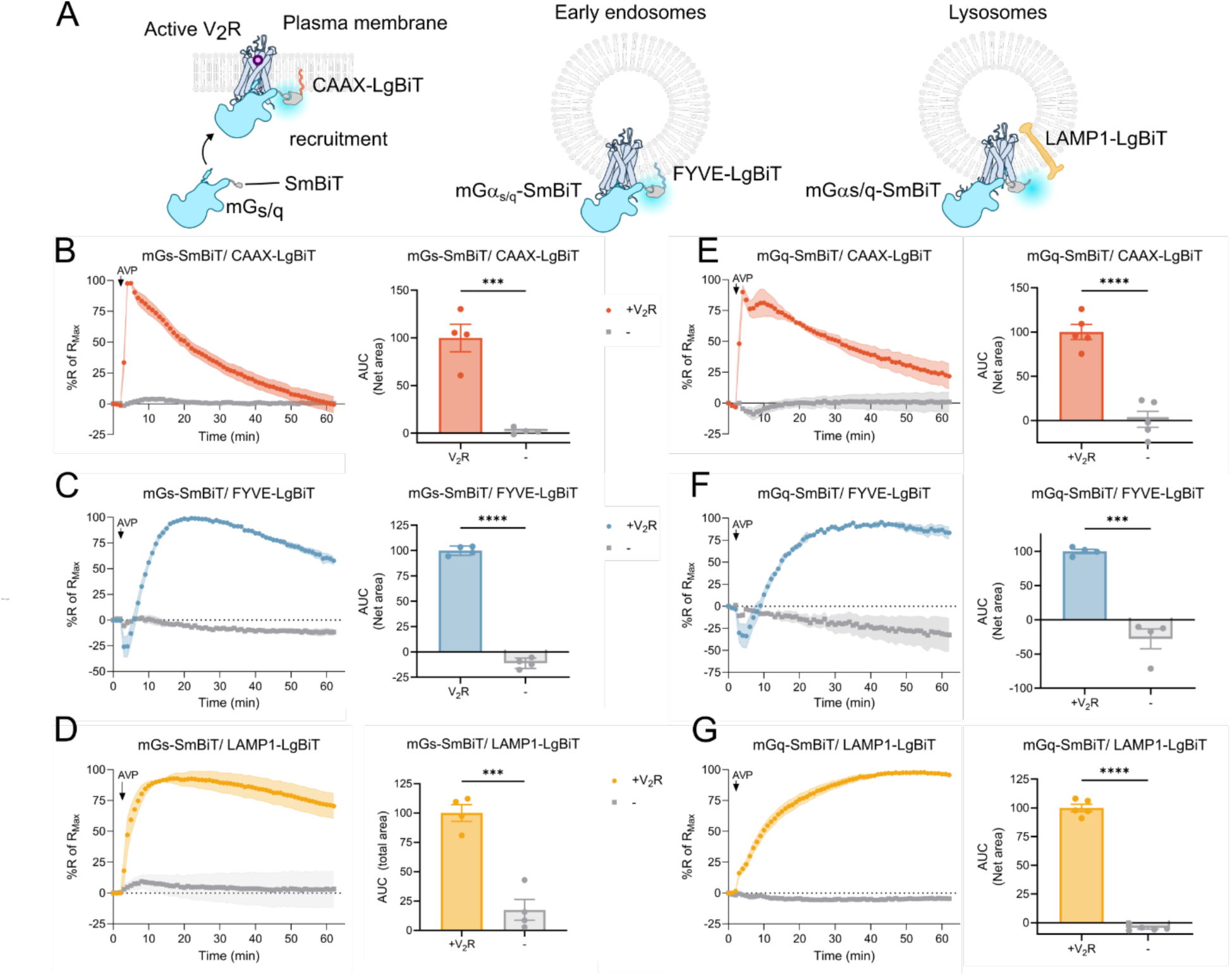
Translocation of mGs and mGq to V2R to the plasma membrane, early endosome, and lysosome in response to V2R activation. **(A)** Schematic representation of how mGs-SmBiT and mGq-SmBiT was used as subrogates of G protein to determine V2R coupling at plasma membrane, early endosomes, and lysosomes. (**B-G**) Time-course experiments measuring AVP-stimulated (100 nM) changes in luminescence signal generated between mGs-SmBiT and CAAX-LgBiT (**B**), FYVE-LgBiT (**C**), or LAMP1-LgBiT (**D**), or between mGq-SmBiT and CAAX-LgBiT (**E**), FYVE-LgBiT (**F**), or LAMP1-LgBiT (**G**). Responses were normalized to the maximal change in luminescence. The experiments were conducted in V2R-expresing HEK293 cells (colored curves) and non-transfected HEK293 cells (gray curves). AUC values were used to calculate the total mGs/q translocation response (bar graphs). Data represent the mean ± SEM from N=4-5 independent experiments. Unpaired t-test with two-tail p-value tests were applied to determine the statistical differences between the distinct treatments (***p < 0.001; ****p < 0.0001).

### V2R activates endogenous G protein and βarr1 at the lysosomal surface

Since overexpression of transducers can trigger cellular responses not observed under endogenous and physiologically relevant conditions, we aimed to investigate whether V2R can activate endogenous G proteins and βarrs. We specifically focused on the ability of V2R to activate endogenous G_s_ and βarr1 at lysosomes.

To examine whether endogenous βarr1 translocate to and is activated at lysosomes, we measured NanoBiT bystander complementation between Ib30-SmBiT and LAMP1-LgBiT in V2R-expressing HEK293 cells (Fig. 4A). Ib30 is an intrabody biosensor that is expressed in the cytosol of mammalian cells and can be used to detect active βarr1 (*41, 42*). The biosensor was derived from the antibody binder fragment Fab30, which selectively recognizes an active conformation of βarr1 when in complex with the phosphorylated C-terminal tail of V2R, and has been applied in structural studies to stabilize GPCR–βarr1 complexes (*42–46*). Upon AVP-stimulation, we observed an increase in the translocation of Ib30-SmBiT to lysosomes in V2R-expressing cells, with kinetics that mirrors that of βarr1-SmBiT (Fig. 4B). This response was not observed in HEK293 cells that do not express the V2R (Fig. 4B). Therefore, these results imply that AVP-activated V2R associates with endogenous and active βarr1 at the lysosomal surface.

**Fig. 4.**
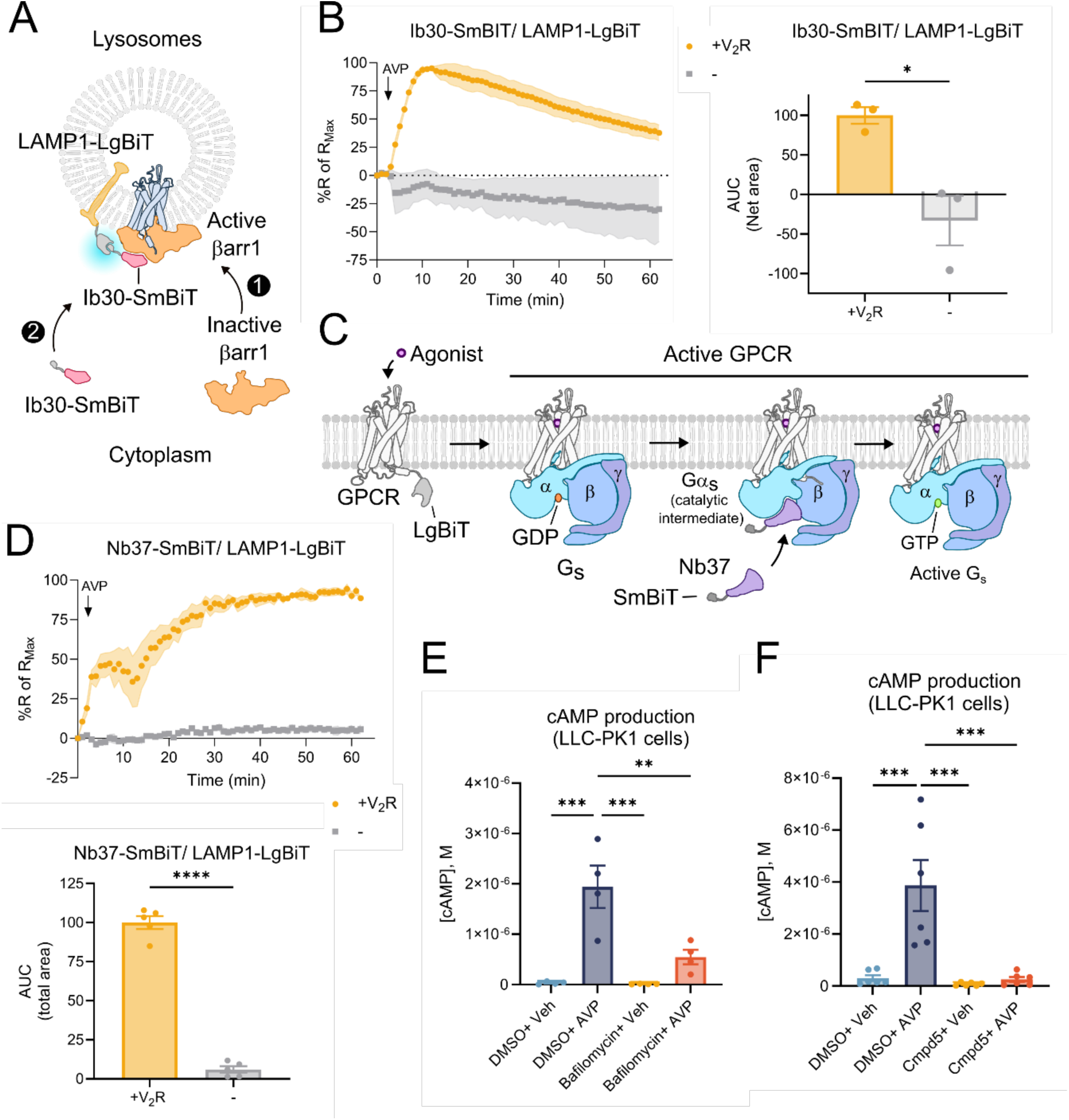
V2R-mediated recruitment and activation of endogenous βarr1 and Gα_s_ at the lysosomal surface and the influence of lysosomal V2R trafficking on total Gs signaling. (**A**) Schematic illustration of how the biosensor Ib30-SmBiT was used in the NanoBiT assay to detect the translocation of active endogenous βarr1 to the lysosomal surface containing LAMP1-LgBiT. (**B**) Time-course experiment measuring AVP-stimulated (100 nM) changes in luminescence as a result of proximity between Ib30-SmBiT and LAMP1-LgBiT in V2R (yellow) or mock-transfected (grey) HEK293 cells. Responses were normalized to the maximal change in luminescence. AUC values were used to calculate the total Ib30-SmBiT/LAMP1-LgBiT response (bar graphs). Data represent the mean ± SEM from N=3 independent experiments. (**C**) Schematic representation of how the Nb37-SmBiT construct is used as a biosensor of active endogenous Gα_s_. (**D**) Real-time NanoBiT assay bystander complementation between Nb37-SmBiT and LAMP1-LgBiT in response to AVP-stimulation (100 nM). The experiments were conducted in V2R-expresing HEK293 cells (yellow curves) and non-transfected HEK293 cells (gray curves). AUC values were used to calculate the total Nb37-SmBiT/LAMP1-LgBiT response (bar graphs). Data represent the mean ± SEM from N=4 independent experiments. (**E**) AVP-stimulated (100 nM) or mock-stimulated cAMP accumulation in LLC-PK1 cells, which express the V2R endogenously. The cells were preincubated for 30 minutes with either bafilomycin A1 (10 µM) or DMSO (0.1%) control. Data represent the mean ± SEM from N=4 independent experiments. (**F**) AVP-stimulated (100 nM) or mock-stimulated cAMP measurements in LLC-PK1 cells, which were preincubated with the negative allosteric βarr1/2 modulator Cmpd-5 (50 µM) or DMSO (0.1%) control. Data represent the mean ± SEM from N=6 independent experiments. (**B** and **D**) Unpaired t-test with two-tail p-value tests were applied to determine statistical differences between the distinct treatments (*p<0.05; ****p < 0.0001). (**E-F**) One-way ANOVA with Tukey-multiple comparison post-test was applied (**p<0.01; ***p < 0.001).

To investigate if the V2R activates endogenous G_s_ at lysosomes, we use the nanobody 37 (Nb37) biosensor that detects active G_s_. Originally, Nb37 was developed to recognize a catalytic intermediate of Gα_s_ activation to stabilize GPCR–G_s_ complexes for structural studies, but was repurposed as a biosensor to detect active G_s_ in mammalian cells (Fig. 4C) (*19, 21, 47*). In wild type HEK293 cells (HEK293 WT) expressing Nb37-SmBiT, LAMP1-LgBiT, and V2R, AVP-stimulation caused an increase in the NanoBiT signal suggesting that G_s_ is being activated at the lysosomal surface (Fig. 4D). This response was not observed in HEK293 cells without V2R expression (Fig. 4D, gray) or in V2R-expressing cells where the Gα_s_ subtypes have been knocked out (HEK293 ΔGα_s_; Supplementary Fig. S3). Thus, these results suggest that AVP-stimulated V2R activates endogenous G_s_ at lysosomes.

To evaluate if lysosomal activity of V2R contributes to downstream signaling events in a physiological-relevant cell model, we used the kidney epithelial cell line LLC-PK1, which expresses the V2R endogenously. As this cell line is challenging to transfect at levels needed for proper expression of our biosensors, we used the bafilomycin A1 to inhibit V2R trafficking to lysosomes and measured how this affects V2R-stimulated G_s_-cAMP signaling. Bafilomycin A1 is a selective small molecule inhibitor of the V-type H^+^-ATPase pump, which uses the free energy harvested from ATP hydrolysis to pump protons into the lumen of lysosomes and other intracellular vesicles (*48*). Inhibition of V-type H^+^-ATPase by bafilomycin A1 disrupts lysosomal physiology and thereby prevents the transport of cholesterol and proteins to the lysosomes (*49, 50*). In fact, pre-incubating HEK293 cells with bafilomycin A1, significantly reduced AVP-induced V2R-SmBiT trafficking to lysosomes as well as V2R-mediated βarr1-SmBiT and mGs-SmBiT recruitment to lysosomes (Supplementary Fig. S4A-C). Interestingly, pre-incubating LLC-PK1 cells with bafilomycin A1, led to a significant reduction in AVP-stimulated cAMP accumulation as compared to DMSO control (Fig. 4E). These results suggest that disruption of cargo trafficking to lysosomes blunts V2R-mediated G_s_ signaling.

As recruitment of βarrs plays a major role in V2R internalization and trafficking to lysosomes, we blocked βarr function in LLC-PK1 cells by pre-incubation with the newly discovered βarr negative allosteric modulator, Cmpd-5 (*51*). In LLC-PK1 cells, pre-incubation of Cmpd-5 significantly reduced AVP-stimulated cAMP accumulation compared to cells that were preincubated with DMSO control (Fig. 4F). This observation indicates that βarr-mediated V2R trafficking positively regulates the integrated G_s_ response in LLC-PK1 cells.

Together, these results indirectly suggest that G_s_ activation at lysosomes by βarr-internalized V2R, contributes to the global cAMP production response in an endogenous and physiologically relevant cell system (Fig. 5).

**Figure 5.**
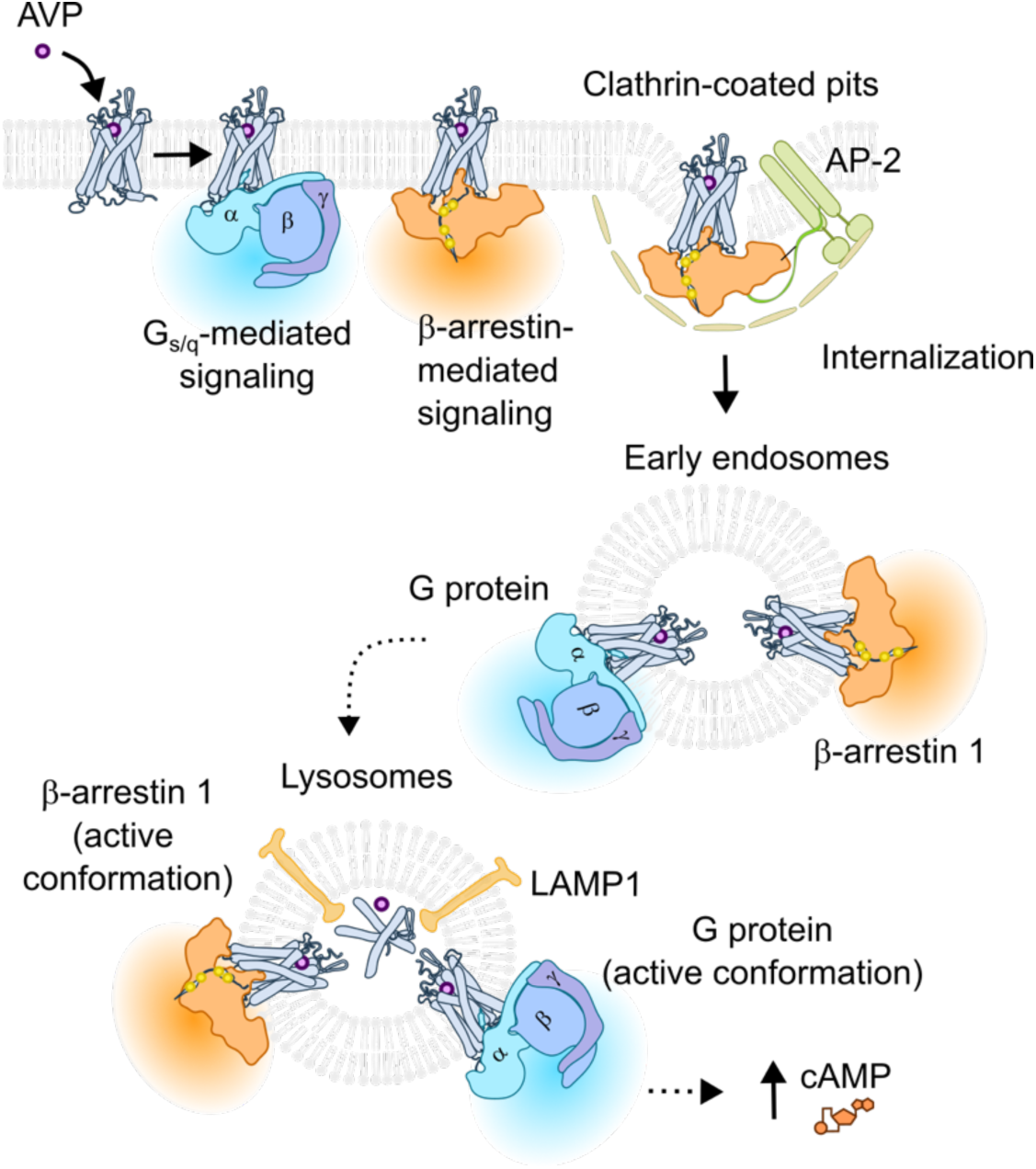
Proposed model for V2R activity at the lysosomal surface. Binding of AVP activates the V2R, which leads to recruitment and activation of G proteins (G_s_ and G_q/11_). At the same time, the V2R is phosphorylated by GRKs and βarrs are recruited to the phosphorylated receptor at the plasma membrane. Next, stable V2R-βarr complexes at clathrin-coated pits are endocytosed into the early endosomes where the receptor retains its ability to bind transducers and promote signaling. Once the receptor is sorted to lysosomes, it still couple to and activate Gα_s_, Gα_q_ and βarr1 at the lysosomal surface before eventually being engulfed into the lumen for degradation.

## Discussion

GPCR trafficking to lysosomes has traditionally been considered a slow process and the final destination of internalized receptors committed to degradation (*52–54*). Despite previous studies reporting on V2R trafficking to lysosomes in cells exposed to AVP for prolonged periods of time (*10, 36*), we here report on a relatively rapid increase in V2R localization at the lysosomal membrane within the first 10-15 minutes of agonist stimulation (Fig. 1D). This lysosomal presence remained relatively stable for up to an hour. We used LAMP1-LgBiT as a lysosomal marker in our NanoBiT approach, a high-throughput technology that permits tracking of cellular protein-protein proximity with high sensitivity in real-time. LAMP1 is one of the most abundant proteins in the lysosome (*55*) and has consistently been used as a lysosomal marker, not only for the studies of lysosomal biology in general, but more importantly, for the trafficking of the V2R to lysosomes (*10, 56*). We verified that the construct co-localizes nearly perfectly with a commercial lysosomal marker that has been extensively used previously (Supplementary Fig. S1A-B) (*57–59*). Moreover, our construct contains the LgBiT part fused to the C-terminal region of LAMP1, which is exposed on the cytosolic side of the lysosome. Therefore, the LAMP1-LgBiT is an ideal construct to measure trafficking of SmBiT-tagged V2R (to the C-terminal tail of the receptor) as well as different transducers to the cytosolic lysosomal surface in real-time.

GPCRs and other membrane proteins are typically sorted to lysosomes for degradation, which makes the potential role of active V2R at the lysosomal surface somewhat puzzling. However, the degradation of V2R requires the receptor to be engulfed into the lysosomal lumen and takes place at a relatively slow pace (*10, 60*). For example, it has been shown in LLC-PK1 cells that prolonged AVP-exposure leads to V2R trafficking to lysosomes, but that only a minor fraction of the receptors is degraded after 1-2 hours of stimulation (*60*). In these cells it takes approximately 4 hours of stimulation to degrade ∼50% of the V2R and up to 24 hours for complete lysosomal degradation (*10, 56*). Our results suggest that the V2R is trafficked much faster to the lysosomal membrane (10-15 minutes) (Fig. 1D). We also showed that V2R at the surface of lysosomes preserved its ability to recruit G proteins and βarrs (Fig. 2-4). In fact, the V2R can bind βarrs exclusively through the C-terminal tail while coupling to G protein via the receptor core, which leads to simultaneous activation of both transducers (*11–14*). Interestingly, a recent study found that a large pool of endogenous G proteins exists at the lysosomal surface (*35*). Therefore, local environment and the biochemical nature of the receptor leave the V2R with plenty of time and resources to promote transducer signaling from this compartment, and thus, lysosomes may represent an unappreciated GPCR signaling hub.

To test whether the V2R activates endogenous G_s_ and βarr1 at the lysosomal surface, we used the biosensors Nb37 and Ib30, respectively (Fig. 4). Nb37 was originally developed for structural studies of the β_2_AR-G_s_ complex (*47*), but was later repurposed as a biosensor that detects Gα_s_ activation by confocal imaging in live cells (*19*). In addition to β_2_AR-activated Gα_s_, Nb37 has been used in a NanoBiT setup to trace Gα_s_ activation by other receptors such as the vasoactive intestinal peptide receptor 1 and glucagon-like peptide-1 receptor (*21, 61*). In those studies, overexpression of G protein subunits (αβγ), either as tricistronic (*21*) or independent plasmids (*61*), was necessary to obtain detectable luminescence signals. Here, we performed our Nb37-SmBiT/LAMP1-LgBiT NanoBiT bystander experiments without co-expression of any G protein subunit, and thus, our data reports activation of *endogenous* G_s_ in real-time at lysosomes. As with Nb37, Ib30 was originally developed as an antibody fragment (Fab30) that recognizes an active βarr1 conformation in complex with the V2Rpp (V2R C-terminal phosphopeptide), and was used for structural studies (*43, 44*). Later, this construct too was repurposed to detect active endogenous βarr1 in live cells when coupled to different Class B GPCRs including the V2R, neurotensin receptor 1, muscarinic M2 receptor, complement C_5a_ receptor, and atypical chemokine receptor ACKR3 (*41, 42*). Using the Ib30-SmBiT, we here showed that the AVP-stimulated V2R complexes with endogenous active βarr1 via its C-terminal tail at the lysosomal surface (Fig. 4B).

Disrupting the acidification in lysosomes and other acidic organelles (endosomes) using the V-type ATPase inhibitor bafilomycin A1, reduced the trafficking of active and signaling competent V2R to lysosomes in HEK293 cells (Supplementary Fig. S4A). Interestingly, bafilomycin A1 treatment reduced AVP-stimulated cAMP production in physiologically relevant LLC-PK1 kidney epithelial cells (Fig. 4E). Moreover, treating LLC-PK1 with a newly discovered βarr negative modulator, Cmpd-5, which inhibit AVP-stimulated βarr recruitment and internalization of V2R (*51*), also reduced AVP-stimulated cAMP production in LLC-PK1 cells (Fig. 4F). These results indirectly suggest that the G_s_-mediated cAMP response depends on functional endolysosomal compartments and βarr-dependent V2R internalization in LLC-PK1 cells. In support of this, bafilomycin A1 has previously been shown to disrupt AVP-induced recycling of AQP2-containing vesicles to the plasma membrane in LLC-PK1 cells (*62*). Moreover, chronic administration of chloroquine, which also disrupts the acidification of intracellular vesicles (*63*), is known to cause polyuria by suppressing the antidiuretic response to vasopressin in the kidney collecting duct (*64*). Interestingly, it was shown that daily injection of chloroquine for 4 days in Sprague-Dawley rats led to polyuria with lower levels of cAMP in the collecting duct. Furthermore, AVP-responsive proteins including AQP2 and the Na^+^-K^+^-2Cl^-^ co-transporter were downregulated as well (*65*).

A conceptual concern of our model arises from the harsh conditions within the lysosomal lumen characterized by an abundance of hydrolases and low pH, which pose a challenge for ligand binding to active GPCRs. Interestingly though, the V2R has evolved to maintain its ability to bind AVP under such harsh conditions as it is expressed in the principal cells along the entire kidney collecting duct, which spans from the cortex to the tip of the papilla (*66*). The pH and osmolarity of this region are known to vary drastically from neutral pH at 7.4 and isosmotic conditions at the cortical interstitium to acidic pH at >5.0 and hyperosmolarity at the tip of the papilla (*67*). Although the affinity for AVP is lower at pH 5.5 than at pH 7.4, the V2R maintain its maximal binding capacity as well as its ability to stimulate G_s_ and internalize under acidic conditions (*68*). In fact, it has been shown that AVP can bind to the V2R in a functional manner at pH as low as 3 (*69*), and thus, the AVP-V2R complex is likely to remain stable throughout its trafficking to the lysosomes. These acid-resistant features of AVP binding to the V2R appear to be unique as another endogenous V2R ligand, oxytocin, does not readily bind the internalized receptor for long periods (*15, 70*). This also explains why V2R stimulation with AVP, but not oxytocin, promotes sustained G_s_ signaling by internalized receptors, which is associated with AQP2 phosphorylation and enhanced ENaC activity at the apical membrane of kidney collecting duct cells (*15*).

Lysosomal signaling may represent a new avenue of GPCR research, not only restricted to Class B receptors, but also on a more general scale. For example, proton-sensing GPCRs (e.g. GPR4, GPR65, GPR68, and GPR132 (*71*)) should in principle be stimulated by acidic conditions of the lysosomal lumen and thereby maintain high signaling capacity after they have been sorted to this compartment. GPR65 has been shown to internalize and constitutively drive G protein signaling from the slightly acidic endosomal compartments (*72*), and may be even more active in the more acidic lysosomal environment. Interestingly, a coding variant of GPR65 (I123L) displayed reduced signaling capacity and is linked to increased risk of inflammatory bowel disease (IBD) (*73*). In IBD patient cells and epithelial cells expressing GPR65 I231L, abnormal lysosomal pH has been observed that causes lysosomal dysfunction and impaired ability to clear bacteria through autophagy (*73*). These findings highlight a role of GPR65 in regulating lysosomal biology potentially via lysosomal signaling. If so, lysosomal signaling may represent a broader mechanism applied by several GPCRs and should be considered in future functional studies and drug discovery efforts.

## Materials and Methods

### Reagents

Dulbecco’s Modified Eagle’s Medium (DMEM) +GlutaMAX-1 containing 4.5 g/L D-Glucose (Cat. 10566-016), Opti-MEM reduced serum/no phenol red (Cat. 11058-021), fetal bovine serum (FBS) premium (Cat. A5670801), penicillin/streptomycin (P/S; Cat. 15140-122), and 0.05% Trypsin-EDTA (Ethylenediaminetetraacetic acid; Cat. 25300-054) were all purchased from Fisher Scientific. Poly-D-lysine (Cat. 150175) was acquired from MP Biomedicals. Lipofectamine 3000 (Cat. L3000015), donkey anti-mouse IgG antibody conjugated to Alexa Fluor 488 (cat. no. A21202), Paraformaldehyde (Cat. 158127), Bafilomycin A1 (Cat. J61835), CellLight Lysosomes-RFP, BacMam 2.0 (Cat. C10597), CellLight Early endosomes-RFP, BacMam 2.0 (Cat. C10587), and Goat anti-Mouse (H+L) Cross-Adsorbed Secondary Antibody, Alexa Fluor™ 488 (Cat. A-11001) were all purchased from Thermo Fisher Scientific. Hank’s balanced salt solution (HBSS), forskolin, HEPES, NaCl, phosphate saline buffer (PBS) tablets, bovine serum albumin (powder isolated), 3-isobutyl-1-methylxanthine (IBMX), KCl, MgCl_2_, NaHCO_3_, NaH_2_PO_4_, glucose, ethylenediaminetetraacetic acid (EDTA), and anti-FLAG M2-HRP conjugated antibody (Cat. A8592, Sigma-Aldrich) were all purchased from Sigma-Aldrich. Coelenterazine h (Cat#301) was purchased from NanoLight Technology. (Arg8)-Vasopressin (AVP; Cat. AB120175) and normal horse serum (Cat. AB7484) were purchased from Abcam. HA-probe mouse monoclonal antibody F-7 (Cat. sc-7392) was purchased from Santa Cruz Biotechnology. Halo Tag ligand 618 (Cat. G980A) was purchased from Promega. SNAP-Surface Alexa Fluor 488 was purchased from New England Biolabs (Cat. S9129S). HTRF cAMP G_s_ Dynamic Kit (Cat. 62AM4PEB) was purchased from Revvity-Cisbio.

### Plasmid constructs

HA-V2R plasmid was N-terminal tagged with an HA-tag (YPYDVPDYA) and cloned and synthesized into pcDNA3.1(+) expression plasmid by GenScript. SNAP-V2R construct was previously described and validated (*14*). βarr1-SmBiT, CAAX-LgBiT, and FYVE-LgBiT plasmids have previously been described (*31, 74*). The LgBiT sequence in those constructs was attached upstream to the sequence coding for CAAX and FYVE. However, we named them CAAX-LgBiT and FYVE-LgBiT along with the text for practical purposes. LAMP1-LgBiT-HA (named LAMP1-LgBiT) construct was designed with the LgBiT sequence fused to the C-terminus of LAMP1 through a flexible linker (GGSGGGGSGGSSSGG), followed by an HA-tag fused downstream to LgBiT, separated with a GGSG linker. mGs and mGq constructs have been described before (*31*). Briefly, mGs and mGq contain the SmBiT sequence fused to the N-terminus of the protein through a GGSG linker, preceded in the N-terminus by a nuclear export signal (MLQNELALKLAGLDINKT) via a GGSG linker to enhance their basal localization in the cytoplasm. Ib30-SmBiT was designed with the SmBiT sequence (MVTGYRLFEEIL) linked to the C-terminus portion of Ib30 through a GGSGGGGSGGSSSGG sequence. SmBiT-Nb37-HA construct (named here Nb37-SmBiT for practical proposes) contains the SmBiT sequence fused to the N-terminus of Nb37 via GGSGGGGSGGSSSGG linker, and an HA-tag fused to the Nb37 C-terminus. Ib30-SmBiT is composed by SmBiT attached to the C-terminus of Ib30, both sequences separated by the same linker used for Nb37. The βarr1-HaloTag (βarr1-HT) was designed with a HaloTag sequence at the C-terminus of βarr1. Nb37-SmBiT, Ib30-SmBiT, V2R-SmBiT, and βarr1-HT constructs were synthesized into the pTwist-CMV vector by Twister Bioscience.

### Cell culture

Parental and Δβarr1/2 (double βarr1 and βarr2 knock-out) HEK293 cells were grown in DMEM media supplemented with 10% fetal bovine serum (FBS) and 1% penicillin/streptomycin (P/S; 100 U/ml), at 37 °C and 5% CO2 incubation conditions. LLC-PK1 cells, a line with epithelial morphology derived from the kidney of a young male pig, were cultured in M199 media supplemented with 3% FBS and 1% P/S at 37 °C, 5% CO2. Cell cultures were passaged every 3– 4 days using trypsin–EDTA 0.05%. For experiments, cultures with up to 15 times passages were used.

### Transfection

Transfections of DNA constructs were performed on HEK293 parental and Δβarr1/2 cells using Lipofectamine 3000. A ratio of 1:5 of the LgBiT/SmBiT DNA constructs, respectively, was used for all the experiments. HA-V2R plasmid or control mock was included in the DNA mix. 1 µL of each Lipofectamine 3000 reagent was diluted in 25 µL of OptiMEM reduced serum medium and added to DNA mixes. Cells cultured in 24-well plates (seeded with 500,000 cells/well a day before transfection) were incubated with the DNA-containing transfection solution for 24 hours. Experiments were performed 48 hours after transfections.

### NanoBiT assay

Bystander NanoBiT assays expressing LgBiT and SmBiT-fused constructs were performed to measure the complementation by shared membrane localization between the proteins. Parental or Δβarr1/2 HEK293 cells were plated on a poly-D-lysine coated 96-well White Flat Bottom Microplate (Falcon^®^, Cat# 353296) and equilibrated in Opti-MEM™ at 37°C for 60 minutes. Coelenterazine-h was added at a final concentration of 10 μM before starting the measurement. After establishing a baseline response for 2 minutes, cells were stimulated with AVP 100 nM and the luminescence was measured for another 60 minutes. In experiments analyzing the effect of lysosomal function inhibition, Bafilomycin A1 10 µM or DMSO 0.1% were added to the cells at the same time as Coelenterazine-h, 30 minutes before adding AVP, with cells being maintained at 37 °C. The signal was detected at 550 nm using a PHERAstar FSX instrument (BMG LabTech). Triplicates of each condition were recorded for the entire time course of AVP stimulation. ΔRLU values were calculated from the RLU values of wells stimulated with AVP minus RLU from wells incubated with vehicle (OptiMEM), both normalized first to the values at time 0. The data presented correspond in each case to the mean of ΔRLU from independent experiments, normalized to the maximal or minimal response in the condition expressing VR.

### HTRF cAMP G_s_ Assay

LLC-PK1 cells preincubated with Bafilomycin A1 10 µM or DMSO 0.1% for 30 minutes were harvested and resuspended in assay buffer (500 nM IBMX + 20 mM HEPES in HBSS buffer, pH 7.4, +Bafilomycin or DMSO at the same concentration) to a final concentration of 2,000,000 cells/mL. In a 384-well Small Volume White Flat Bottom Microplate (Greiner Bio-One GmbH, Cat# 784075), 5 μL of ligand buffer (assay buffer + 200nM AVP or 20 μM forskolin), or vehicle was mixed with 5 μL of the cell suspension (10,000 cells). The plate was incubated at 37°C for 30 minutes before adding 10 μL of lysis buffer containing cAMP Eu-cryptate antibody working solution and cAMP-d2 reagent working solution (HTRF cAMP G_s_ Dynamic Kit). The plate was incubated for additional 1 hour at room temperature. The fluorescence was measured on the PHERAstar *FSX* where wells were excited with light at 340 nm and emission light was measured at 615 nm and 665 nm. The TR-FRET 665 nm/615 nm ratio, which is inversely proportional to the cAMP concentration, was used in combination with a cAMP standard curve to calculate the cAMP production in the cells.

### Confocal microscopy

HEK293 cells transfected with the appropriate constructs (500,000 cells) were transferred to 35 mm glass-bottom Petri dishes (MatTek Cat. PG35G-1.5-10-C) pretreated with poly-D-lysine 24 hours before the experiments. Transduction of HEK293 cells with 10 µL/dish of CellLight Lysosomes-RFPBacMam 2.0 or CellLight Early Endosomes-RFPBacMam 2.0, to label lysosomes with LAMP1-red fluorescent protein (LAMP1-RFP) or early endosomes with Rab5a-RFP, was performed on suspended cells transferred to glass-bottom Petri Dishes. Experiments were performed 24 hours after transduction. For cells transfected with the βarr1-HT construct, HaloTag 618 ligand (100 nM final concentration) was added to the culture media overnight the day before the experiments. For cells where agonist-induced effect was assessed, cultures were incubated with FBS-free DMEM for an hour prior to starting the stimulation with 100 nM AVP or vehicle (FBS-free DMEM) for 30 minutes. For experiments with SNAP-tagged V2R, cells were incubated with FBS-free DMEM for 30 minutes before adding the SNAP-Surface 488 substrate (at a final concentration of 5 µM) for 30 minutes to label V2R localized at the plasma membrane. Stimulation protocol started immediately after washing the SNAP-Surface Alexa Fluor 488 (SNAP488) two times with PBS and adding serum-free media. After the stimulation protocol, with stimulation for 30 minutes with AVP or vehicle, cells were washed two times with PBS and fixed with 4% paraformaldehyde in PBS for 20 minutes on ice, washed and maintained in PBS at 4°C until observation under a confocal microscope. For immunolabelling of LAMP1-LgBiT-HA, cells were incubated with a blocking buffer (PBS supplemented with 0.3% saponin, and 3% normal horse serum) for 60 minutes at room temperature. After that, cells were incubated overnight with the HA-probe mouse antibody (1:200 dilution in blocking buffer) at 4° C. Anti-mouse coupled to Alexa Fluor 488 secondary antibody (1 µg/mL) was incubated for 60 minutes at room temperature. Images were obtained with a Leica SP8 confocal microscope (Leica Microsystems) with a 63x oil objective lens (NA 1.30). In general, 5 to 10 images capturing cells were acquired by dish, representing 3 independent experiments. Obtained images were processed using Fiji (version 2.14.0/1.54f; NIH) (*75*) for linear brightness and contrast adjustment applied equally to the entire image data set and adding scale bars. Co-localization was measured using CellProfiler (version 4.2.6, Broad Institute) (*76*) and the colocalization pipeline available in the developer website in order to obtain Pearson’s colocalization coefficients with an automatized quantification process.

## Data and statistical analysis

Data is presented as the mean ± SEM of independent biological units (experiments). Unpaired t-test with two-tail p-value tests were applied when comparing between two groups. One-way ANOVA with Tukey post-tests were applied when comparing between multiple groups. P-values are indicated as: *p<0.05, **p<0.01, ***p<0.001, and ****p<0.0001. All the statistical analysis and graphic representations were obtained with GraphPad Prism 10.5.0 (GraphPad Software).

## Acknowledgments

We thank Asuka Inoue for providing the HEK293 Δβarr1/2 (double knockout) cells. **Funding:** This work was supported by grants from NIH (R35GM147088 (NIGMS) and R21CA243052 (NCI) to A.R.B.T.) and the LEO Foundation (LF18043 to A.R.B.T). **Author contributions:** A.R.B.T. and E.F.E. designed the project. E.F.E. and H.H. performed experiments. E.F.E. analyzed the data. E.F.E. and A.R.B.T. wrote the manuscript. **Competing interests:** A.R.B.T. is a scientific co-founder of Unco Therapeutics LLC. The other authors declare that they have no competing interests. **Data and materials availability:** All data needed to evaluate the conclusions in the paper are present in the paper or the Supplementary Materials. The plasmids are available from A.R.B.T. under a material transfer agreement with NYU, USA.

## Supplementary figures

**Supplementary Fig. S1.**
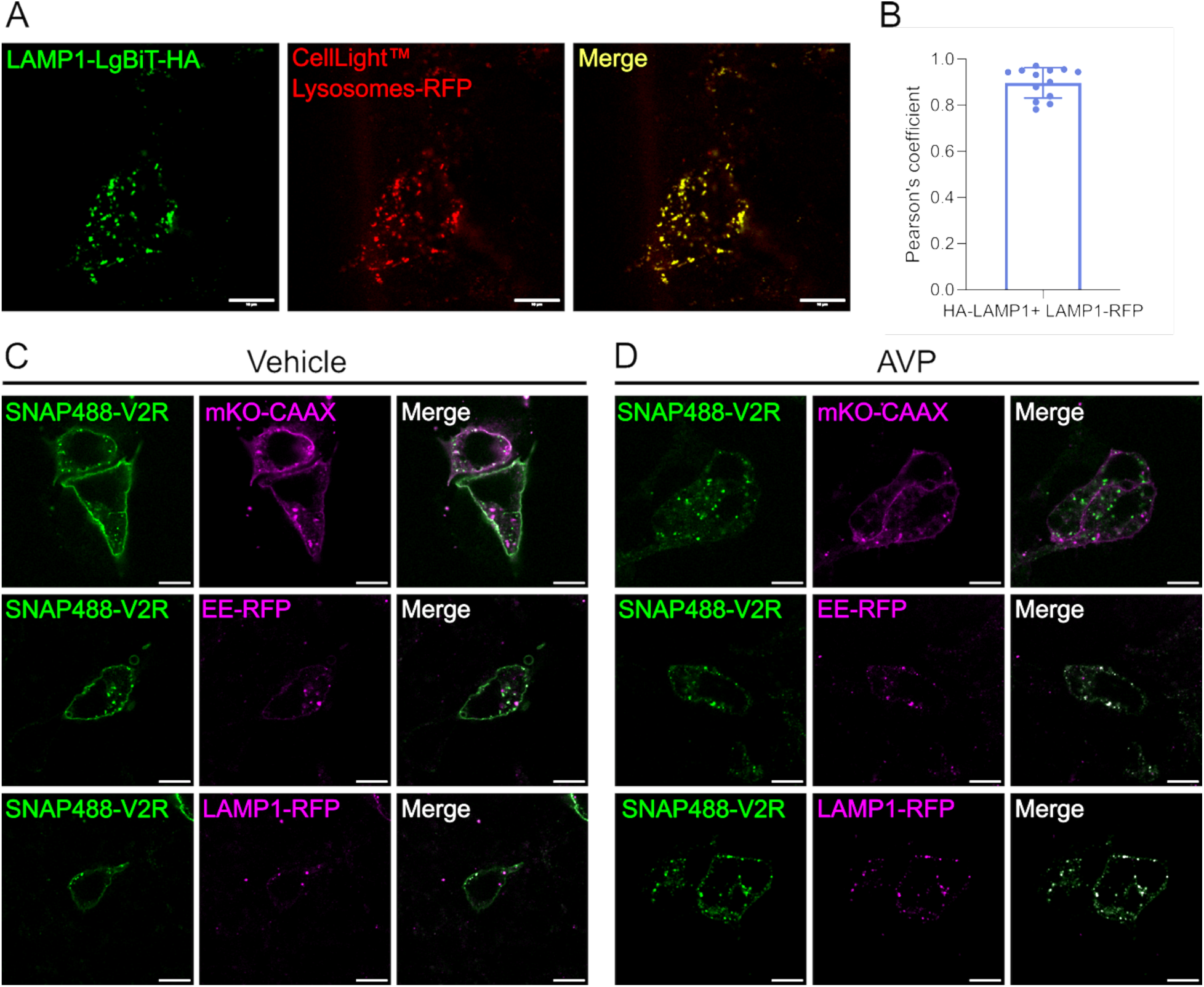
LAMP1-LgBiT colocalization with other LAMP1 marker and SNAP-V2R internalization. (A) Immunofluorescence imaging of HEK293 cells expressing the LAMP1-LgBiT-HA construct (green) and the commercial CellLight™ Lysosomes-RFP marker (red). The cells were stained using a primary anti-HA antibody. Colocalization is observed in white color in the merge image. (B) Quantification of LAMP1-LgBiT-HA and LAMP1-RFP colocalization using Pearson’s colocalization coefficient. Individual values corresponding to cells from 3 independent experiments are shown and the data represents the mean ± SD. (**C-D**) Confocal imagining of HEK293 cells expressing SNAP-V2R construct (green), the plasma membrane marker mKO-CAAX, early endosomes marker EE-RFP, or lysosomal marker LAMP1-RFP (magenta). The cells were stimulated for 30 minutes with vehicle (**C**) or 100 nM AVP (**D**). Merge image shows the composite of the green and magenta panels with white color representing colocalization of the two colors. In all images, the white bar corresponds to the scale of 10 µm.

**Supplementary Fig. S2.**
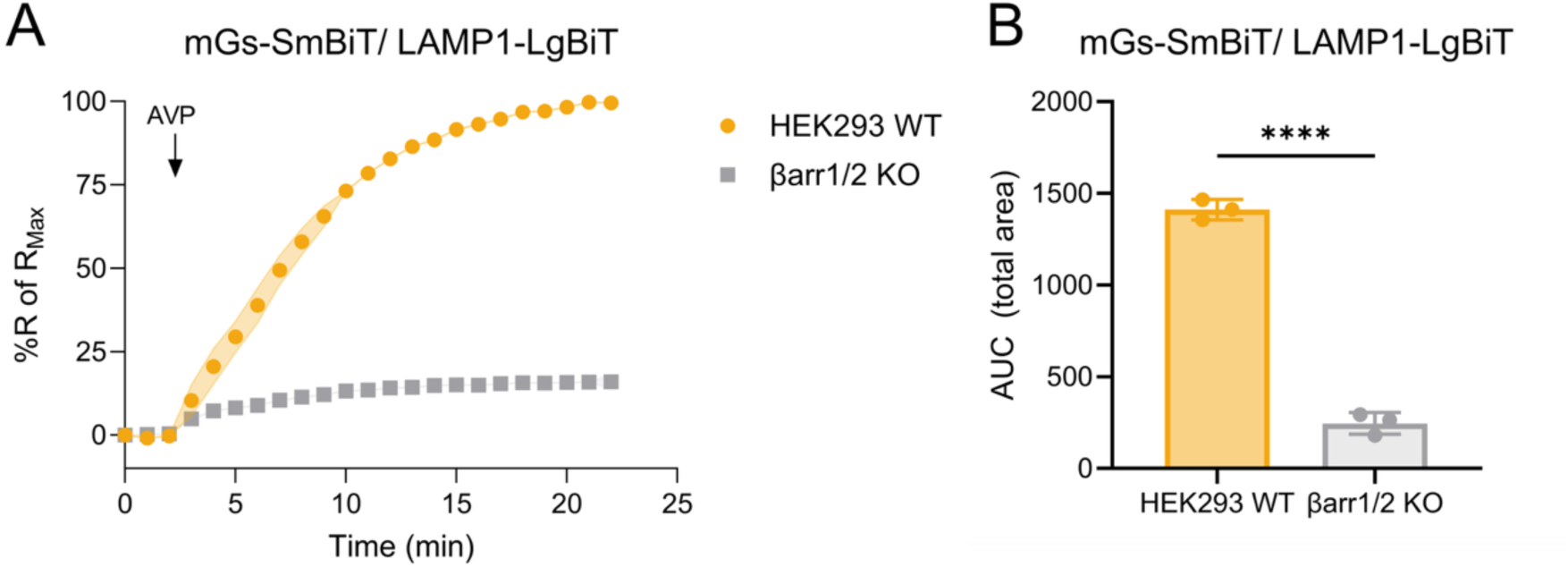
AVP-stimulated mGs-SmBiT recruitment to LAMP1-LgBiT lysosomes in V2R-expressing HEK293 wild-type (WT) or βarr1/2-knockout cells. (A) Time-course NanoBiT experiment detecting mGs-SmBiT and LAMP1-LgBiT proximity in cells stimulated with AVP 100 nM. The experiments were conducted in V2R-expresing HEK293 parental cells (yellow curves) or HEK293 Δβarr1/2 cells (gray curves). Responses were normalized to the maximal change in luminescence. (**B**) AUC values were used to calculate the total mGs-SmBiT/LAMP1-LgBiT response (bar graphs). Data represent the mean ± SEM from N=4 independent experiments.HEK293. Unpaired t-test with two-tail p-value tests were applied to determine statistical differences between the distinct treatments (****p < 0.0001).

**Supplementary Fig. S3.**
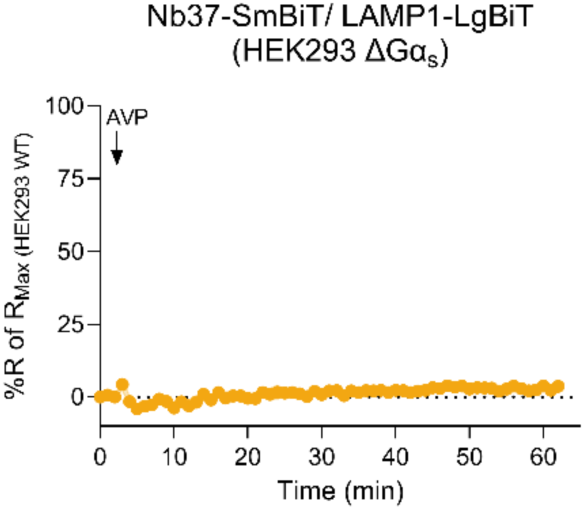
AVP-stimulated Nb37 recruitment to LAMP1-LgBiT in V2R-expressing HEK293 ΔGa_s_ cells. Time-course NanoBiT experiments detecting AVP-stimulated (100 nM) proximity between Nb37-SmBiT and LAMP1-LgBiT in V2R-expressing HEK293 cells where the Gα_s_ has been knocked out. Responses were normalized to the maximal change in luminescence in V2R-expressing HEK293 parental cells. Data represents the mean ± SEM from N=4 independent experiments.

**Supplementary Fig. S4.**
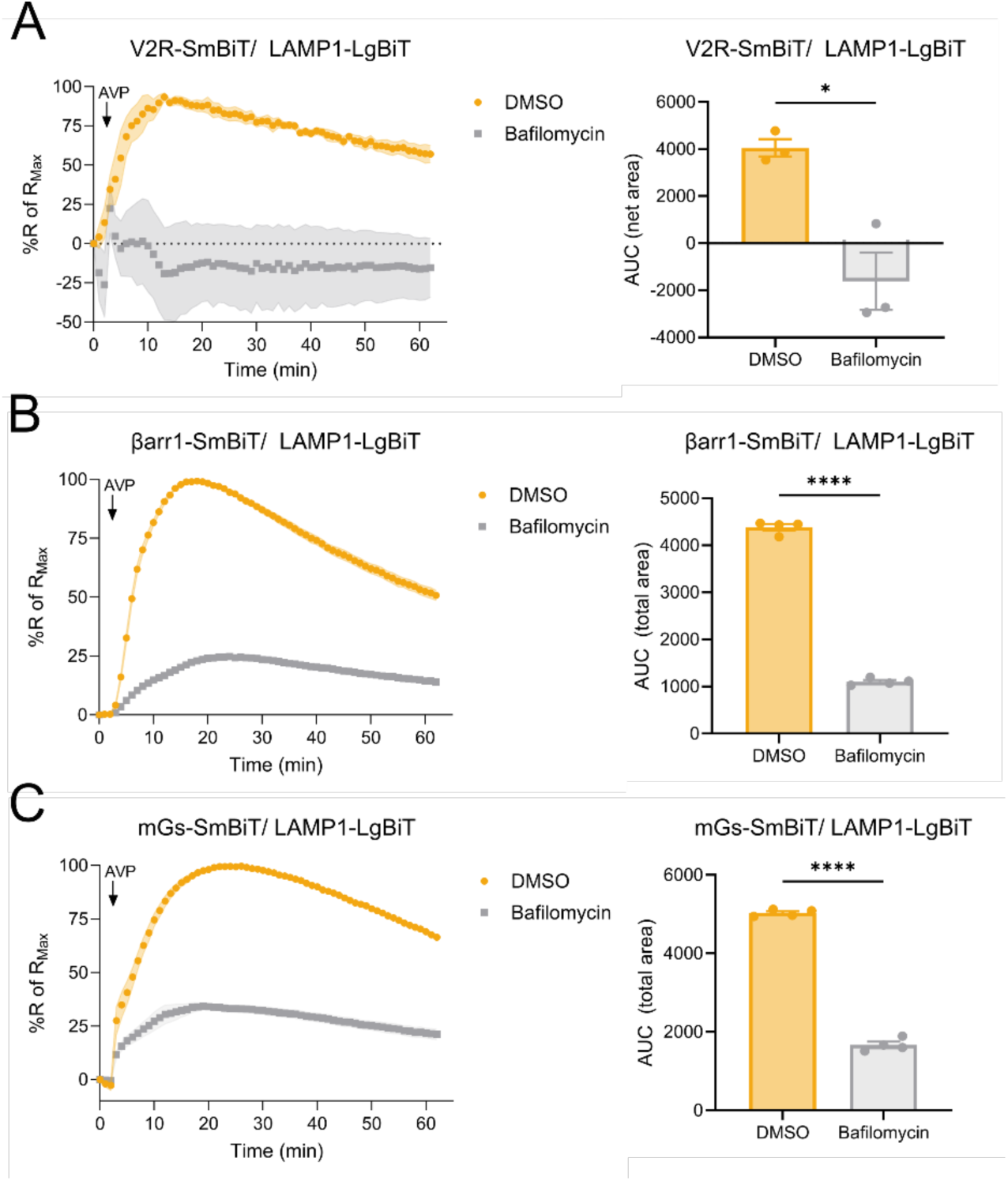
Bafilomycin A1 pretreatment inhibits AVP-induced V2R, βarr1 and mGs localization in lysosomes. (**A**) The effect of 10 µM Bafilomycin (grey) or DMSO control (yellow) pretreatment on AVP-stimulated (100 nM) V2R-SmBiT trafficking to the lysosomal surface in V2R-expressing HEK293 cells. (**B**) AVP-stimulated (100 nM) βarr1-SmBiT recruitment to lysosomes measured as bystander complementation with LAMP1-LgBiT in V2R-expressing HEK293 cells. Cells were pretreated with Bafilomycin (grey) or DMSO control (yellow). (**C**) AVP-stimulated (100 nM) mGs-SmBiT translocation to lysosomes using the NanoBiT approach in V2R-expressing HEK293 cells. Cells were pretreated with Bafilomycin (grey) or DMSO control (yellow). (**A-C**) Responses were normalized to the maximal change in luminescence. AUC values were used to calculate the total translocation response (bar graphs). Data represent the mean ± SEM from N=3-4 independent experiments. Unpaired t-test with two-tail p-value tests were applied to determine statistical differences between the conditions (*p < 0.05; ****p<0.0001).

## References

1. K. V. Juul, D. G. Bichet, S. Nielsen, J. P. Nørgaard, The physiological and pathophysiological functions of renal and extrarenal vasopressin V2 receptors. American Journal of Physiology-Renal Physiology 306, F931–F940 (2014).

2. L. S. Erdélyi, L. Hunyady, A. Balla, V2 vasopressin receptor mutations: future personalized therapy based on individual molecular biology. Frontiers in Endocrinology Volume 14 **-** 2023, (2023).

3. F. Zhou et al., Molecular basis of ligand recognition and activation of human V2 vasopressin receptor. Cell Research 31, 929–931 (2021).

4. H. B. Moeller, J. Praetorius, M. R. Rützler, R. A. Fenton, Phosphorylation of aquaporin-2 regulates its endocytosis and protein–protein interactions. Proceedings of the National Academy of Sciences 107, 424–429 (2010).

5. S. Nielsen, S. R. DiGiovanni, E. I. Christensen, M. A. Knepper, H. W. Harris, Cellular and subcellular immunolocalization of vasopressin-regulated water channel in rat kidney. Proceedings of the National Academy of Sciences 90, 11663–11667 (1993).

6. R. H. Oakley, S. A. Laporte, J. A. Holt, L. S. Barak, M. G. Caron, Association of beta-arrestin with G protein-coupled receptors during clathrin-mediated endocytosis dictates the profile of receptor resensitization. J Biol Chem 274, 32248–32257 (1999).

7. R. H. Oakley, S. A. Laporte, J. A. Holt, M. G. Caron, L. S. Barak, Differential affinities of visual arrestin, beta arrestin1, and beta arrestin2 for G protein-coupled receptors delineate two major classes of receptors. J Biol Chem 275, 17201–17210 (2000).

8. Q. T. He et al., Structural studies of phosphorylation-dependent interactions between the V2R receptor and arrestin-2. Nat Commun 12, 2396 (2021).

9. R. H. Oakley, S. A. Laporte, J. A. Holt, L. S. Barak, M. G. Caron, Molecular determinants underlying the formation of stable intracellular G protein-coupled receptor-beta-arrestin complexes after receptor endocytosis*. J Biol Chem 276, 19452–19460 (2001).

10. R. Bouley et al., Downregulation of the vasopressin type 2 receptor after vasopressin-induced internalization: involvement of a lysosomal degradation pathway. Am J Physiol Cell Physiol 288, C1390–1401 (2005).

11. C. Daly et al., beta-Arrestin-dependent and -independent endosomal G protein activation by the vasopressin type 2 receptor. Elife 12, (2023).

12. T. J. Cahill3rd, et al., Distinct conformations of GPCR-beta-arrestin complexes mediate desensitization, signaling, and endocytosis. Proc Natl Acad Sci U S A 114, 2562–2567 (2017).

13. P. Kumari et al., Core engagement with beta-arrestin is dispensable for agonist-induced vasopressin receptor endocytosis and ERK activation. Mol Biol Cell 28, 1003–1010 (2017).

14. A. R. B. Thomsen et al., GPCR-G Protein-beta-Arrestin Super-Complex Mediates Sustained G Protein Signaling. Cell 166, 907–919 (2016).

15. T. N. Feinstein et al., Noncanonical control of vasopressin receptor type 2 signaling by retromer and arrestin. J Biol Chem 288, 27849–27860 (2013).

16. S. Ferrandon et al., Sustained cyclic AMP production by parathyroid hormone receptor endocytosis. Nat Chem Biol 5, 734–742 (2009).

17. D. Calebiro et al., Persistent cAMP-signals triggered by internalized G-protein-coupled receptors. PLoS Biol 7, e1000172 (2009).

18. E. Flores-Espinoza, A. R. B. Thomsen, Beneath the surface: endosomal GPCR signaling. Trends Biochem Sci 49, 520–531 (2024).

19. R. Irannejad et al., Conformational biosensors reveal GPCR signalling from endosomes. Nature 495, 534–538 (2013).

20. N. M. Fisher, M. von Zastrow, Opioid receptors reveal a discrete cellular mechanism of endosomal G protein activation. Proc Natl Acad Sci U S A 122, e2420623122 (2025).

21. E. E. Blythe, M. von Zastrow, beta-Arrestin-independent endosomal cAMP signaling by a polypeptide hormone GPCR. Nat Chem Biol 20, 323–332 (2024).

22. Y.-J. I. Jong, V. Kumar, K. L. O’Malley, Intracellular Metabotropic Glutamate Receptor 5 (mGluR5) Activates Signaling Cascades Distinct from Cell Surface Counterparts*. Journal of Biological Chemistry 284, 35827–35838 (2009).

23. F. Liccardo et al., Subcellular activation of β-adrenergic receptors using a spatially restricted antagonist. Proceedings of the National Academy of Sciences 121, e2404243121 (2024).

24. A. Godbole, S. Lyga, M. J. Lohse, D. Calebiro, Internalized TSH receptors en route to the TGN induce local Gs-protein signaling and gene transcription. Nature Communications 8, 443 (2017).

25. Y. Suofu et al., Dual role of mitochondria in producing melatonin and driving GPCR signaling to block cytochrome c release. Proceedings of the National Academy of Sciences 114, E7997–E8006 (2017).

26. Q. Wang et al., 5-HTR3 and 5-HTR4 located on the mitochondrial membrane and functionally regulated mitochondrial functions. Scientific Reports 6, 37336 (2016).

27. V. Kumar, Y.-J. I. Jong, K. L. O’Malley, Activated Nuclear Metabotropic Glutamate Receptor mGlu5 Couples to Nuclear G q/11 Proteins to Generate Inositol 1,4,5-Trisphosphate-mediated Nuclear Ca2+ Release *. Journal of Biological Chemistry 283, 14072-14083 (2008).

28. B. Boivin et al., Functional β-adrenergic receptor signalling on nuclear membranes in adult rat and mouse ventricular cardiomyocytes. Cardiovascular Research 71, 69–78 (2006).

29. A. D. White et al., Spatial bias in cAMP generation determines biological responses to PTH type 1 receptor activation. Science Signaling 14, eabc5944 (2021).

30. A. Hegron et al., Therapeutic antagonism of the neurokinin 1 receptor in endosomes provides sustained pain relief. Proceedings of the National Academy of Sciences 120, e2220979120 (2023).

31. H. Hahn et al., Endosomal chemokine receptor signalosomes regulate central mechanisms underlying cell migration. eLife 13, RP99373 (2025).

32. R. E. Yarwood et al., Endosomal signaling of the receptor for calcitonin gene-related peptide mediates pain transmission. Proc Natl Acad Sci U S A 114, 12309–12314 (2017).

33. J. Zhao, S. Benlekbir, J. L. Rubinstein, Electron cryomicroscopy observation of rotational states in a eukaryotic V-ATPase. Nature 521, 241–245 (2015).

34. M. Cao, X. Luo, K. Wu, X. He, Targeting lysosomes in human disease: from basic research to clinical applications. Signal Transduction and Targeted Therapy 6, 379 (2021).

35. W. Jang, K. Senarath, G. Feinberg, S. Lu, N. A. Lambert, Visualization of endogenous G proteins on endosomes and other organelles. Elife 13, (2024).

36. N. P. Martin, R. J. Lefkowitz, S. K. Shenoy, Regulation of V2 vasopressin receptor degradation by agonist-promoted ubiquitination. J Biol Chem 278, 45954–45959 (2003).

37. E. V. Moo, J. R. van Senten, H. Bräuner-Osborne, T. C. Møller, Arrestin-Dependent and - Independent Internalization of G Protein–Coupled Receptors: Methods, Mechanisms, and Implications on Cell Signaling. Molecular Pharmacology 99, 242–255 (2021).

38. B. Carpenter, C. G. Tate, Engineering a minimal G protein to facilitate crystallisation of G protein-coupled receptors in their active conformation. Protein Eng Des Sel 29, 583–594 (2016).

39. R. Nehme et al., Mini-G proteins: Novel tools for studying GPCRs in their active conformation. PLoS One 12, e0175642 (2017).

40. Q. Wan et al., Mini G protein probes for active G protein–coupled receptors (GPCRs) in live cells. Journal of Biological Chemistry 293, 7466–7473 (2018).

41. M. Baidya et al., Allosteric modulation of GPCR-induced beta-arrestin trafficking and signaling by a synthetic intrabody. Nat Commun 13, 4634 (2022).

42. M. Baidya et al., Genetically encoded intrabody sensors report the interaction and trafficking of beta-arrestin 1 upon activation of G-protein-coupled receptors. J Biol Chem 295, 10153–10167 (2020).

43. A. K. Shukla et al., Structure of active beta-arrestin-1 bound to a G-protein-coupled receptor phosphopeptide. Nature 497, 137–141 (2013).

44. A. K. Shukla et al., Visualization of arrestin recruitment by a G-protein-coupled receptor. Nature 512, 218–222 (2014).

45. Y. Lee et al., Molecular basis of beta-arrestin coupling to formoterol-bound beta(1)-adrenoceptor. Nature 583, 862–866 (2020).

46. C. Cao et al., Signaling snapshots of a serotonin receptor activated by the prototypical psychedelic LSD. Neuron 110, 3154–3167 e3157 (2022).

47. G. H. Westfield et al., Structural flexibility of the G alpha s alpha-helical domain in the beta2-adrenoceptor Gs complex. Proc Natl Acad Sci U S A 108, 16086–16091 (2011).

48. T. Yoshimori, A. Yamamoto, Y. Moriyama, M. Futai, Y. Tashiro, Bafilomycin A1, a specific inhibitor of vacuolar-type H(+)-ATPase, inhibits acidification and protein degradation in lysosomes of cultured cells. Journal of Biological Chemistry 266, 17707–17712 (1991).

49. N. Matsumoto, M. Nakanishi-Matsui, Proton pumping V-ATPase inhibitor bafilomycin A1 affects Rab7 lysosomal localization and abolishes anterograde trafficking of osteoclast secretory lysosomes. Biochemical and Biophysical Research Communications 510, 421–426 (2019).

50. T. Furuchi, K. Aikawa, H. Arai, K. Inoue, Bafilomycin A1, a specific inhibitor of vacuolar-type H(+)-ATPase, blocks lysosomal cholesterol trafficking in macrophages. Journal of Biological Chemistry 268, 27345–27348 (1993).

51. A. W. Kahsai et al., Small-molecule modulation of β-arrestins. bioRxiv, 2024.2012.2027.630464 (2025).

52. A. Marchese, M. M. Paing, B. R. S. Temple, J. Trejo, G Protein–Coupled Receptor Sorting to Endosomes and Lysosomes. Annual Review of Pharmacology and Toxicology 48, 601–629 (2008).

53. S. K. Shenoy, P. H. McDonald, T. A. Kohout, R. J. Lefkowitz, Regulation of Receptor Fate by Ubiquitination of Activated &#x3b2; 2-Adrenergic Receptor and &#x3b2;-Arrestin. Science 294, 1307–1313 (2001).

54. M. R. Dores, J. Trejo, Endo-lysosomal sorting of G-protein-coupled receptors by ubiquitin: Diverse pathways for G-protein-coupled receptor destruction and beyond. Traffic 20, 101–109 (2019).

55. E.-L. Eskelinen, Roles of LAMP-1 and LAMP-2 in lysosome biogenesis and autophagy. Molecular Aspects of Medicine 27, 495–502 (2006).

56. S. Chen et al., Visualizing microtubule-dependent vasopressin type 2 receptor trafficking using a new high-affinity fluorescent vasopressin ligand. Endocrinology 152, 3893–3904 (2011).

57. A. B. Dyve Lingelem, J. Bergan, K. Sandvig, Inhibitors of Intravesicular Acidification Protect Against Shiga Toxin in a pH-Independent Manner. Traffic 13, 443–454 (2012).

58. R. S. Ames, T. A. Kost, J. P. Condreay, BacMam technology and its application to drug discovery. Expert Opinion on Drug Discovery 2, 1669–1681 (2007).

59. Y. Yan et al., Particles on the Move: Intracellular Trafficking and Asymmetric Mitotic Partitioning of Nanoporous Polymer Particles. ACS Nano 7, 5558–5567 (2013).

60. X. Yi et al., Alix (AIP1) is a vasopressin receptor (V2R)-interacting protein that increases lysosomal degradation of the V2R. Am J Physiol Renal Physiol 292, F1303–1313 (2007).

61. A. I. Oqua et al., Molecular mapping and functional validation of GLP-1R cholesterol binding sites in pancreatic beta cells. eLife 13, RP101011 (2025).

62. C. E. Gustafson et al., Recycling of AQP2 occurs through a temperature- and bafilomycin-sensitive trans-Golgi-associated compartment. American Journal of Physiology-Renal Physiology 278, F317–F326 (2000).

63. M. A. A. Al-Bari, Targeting endosomal acidification by chloroquine analogs as a promising strategy for the treatment of emerging viral diseases. Pharmacology Research & Perspectives 5, e00293 (2017).

64. M. H. Ahmed, N. Ashton, R. J. Balment, Renal function in a rat model of analgesic nephropathy: effect of chloroquine. The Journal of pharmacology and experimental therapeutics 305, 123–130 (2003).

65. T. N. von Bergen, M. A. Blount, Chronic use of chloroquine disrupts the urine concentration mechanism by lowering cAMP levels in the inner medulla. American Journal of Physiology-Renal Physiology 303, F900–F905 (2012).

66. R. M. Edwards, W. Trizna, L. B. Kinter, Renal microvascular effects of vasopressin and vasopressin antagonists. American Journal of Physiology-Renal Physiology 256, F274–F278 (1989).

67. E. A. Alexander, J. H. Schwartz, Regulation of Acidification in the Rat Inner Medullary Collecting Duct. American Journal of Kidney Diseases 18, 612–618 (1991).

68. E. A. Zalyapin et al., Effects of the renal medullary pH and ionic environment on vasopressin binding and signaling. Kidney International 74, 1557–1567 (2008).

69. D. A. Jans, R. Peters, P. Jans, F. Fahrenholz, Vasopressin V2-receptor mobile fraction and ligand-dependent adenylate cyclase activity are directly correlated in LLC-PK1 renal epithelial cells. Journal of Cell Biology 114, 53–60 (1991).

70. L. B. Teixeira et al., Sustained Galpha(s) signaling mediated by vasopressin type 2 receptors is ligand dependent but endocytosis and beta-arrestin independent. Sci Signal 18, eadf6206 (2025).

71. M. Sisignano, M. J. M. Fischer, G. Geisslinger, Proton-Sensing GPCRs in Health and Disease. Cells 10, 2050 (2021).

72. L. M. Morales Rodriguez, S. E. Crilly, J. B. Rowe, D. G. Isom, M. A. Puthenveedu, Location-biased activation of the proton-sensor GPR65 is uncoupled from receptor trafficking. Proc Natl Acad Sci U S A 120, e2302823120 (2023).

73. K. G. Lassen et al., Genetic Coding Variant in GPR65 Alters Lysosomal pH and Links Lysosomal Dysfunction with Colitis Risk. Immunity 44, 1392–1405 (2016).

74. R. Latorre et al., Mice expressing fluorescent PAR 2 reveal that endocytosis mediates colonic inflammation and pain. Proceedings of the National Academy of Sciences 119, e2112059119 (2022).

75. J. Schindelin et al., Fiji: an open-source platform for biological-image analysis. Nature Methods 9, 676–682 (2012).

76. D. R. Stirling et al., CellProfiler 4: improvements in speed, utility and usability. BMC Bioinformatics 22, 433 (2021).

